# Local climate change velocities explain multidirectional range shifts in a North American butterfly assemblage

**DOI:** 10.1101/2023.07.31.551397

**Authors:** Carmen R.B. da Silva, Sarah E. Diamond

## Abstract

Species are often expected to shift their distributions poleward to evade warming climates. However, from 18 years of fixed transect monitoring data on 88 species of butterfly in the midwestern United States, we show that butterflies are shifting their centroids in all directions, except towards the region that is warming the fastest (southeast). Butterflies shifted their centroids at a mean rate of 4.87 km yr^-1^. The rate of centroid shift was significantly associated with local climate change velocity (temperature and precipitation), but not with mean climate change velocity throughout the species’ ranges. Surprisingly, the centroid shift was also unrelated to species traits expected to mediate the shift response including thermal niche breadth (range of climates butterflies experience throughout their distribution) and wingspan (often used as metric for dispersal capability). Contrasting with a number of previous studies, we observed relatively high phylogenetic signal in the rate and direction species shifted their centroids, suggesting that evolutionary history helps to explain multidirectional range shift responses and that some groups of species will be better able to shift their ranges than others. This research shows important signatures of multidirectional range shifts (latitudinal and longitudinal) and uniquely shows that local climate change velocities are more important in driving range shifts than the mean climate change velocity throughout a species’ entire range.

## Introduction

A majority of studies that have examined terrestrial geographic range shift responses to climate change have focused on poleward and upslope range shift responses (Lenoir et al., 2020; Lenoir & Svenning, 2015; VanDerWal et al., 2013). This is because temperature is correlated with both latitude and elevation, and species are expected to move towards cooler higher latitude and elevation environments with further warming (Chen et al., 2011; Parmesan & Yohe, 2003). However, this might limit our understanding of organismal range shifts as climatic factors are also changing across the longitudinal axis, particularly precipitation (Hordley et al., 2023; Lenoir & Svenning, 2015; VanDerWal et al., 2013). There is accumulating evidence of strong longitudinal components to species range shifts, especially within Lepidoptera and birds (Hordley et al., 2023; Huang et al., 2017, 2023; VanDerWal et al., 2013). While it is not entirely clear what drives these longitudinal shifts, they could arise from a number of non-climatic factors such as biotic interactions and changes in land-use (Caro-Miralles & Gutiérrez, 2023; Kwon et al., 2021; Lenoir et al., 2020; Zhang et al., 2019).

Another possibility is that organisms are responding to changes in climate, but the magnitude and direction of climatic shifts is not strictly in a poleward or upslope direction (Lenoir et al., 2010). There are examples of this phenomenon in mountain plants where the direction of the isotherm or isohyet is downslope and plants are moving downslope accordingly (Crimmins et al., 2011; Lenoir et al., 2010, 2020). In addition, rates of climate change are not uniform across space and species could be shifting away from regions that are changing rapidly, even if that means shifting in a non-poleward or upslope direction (Pinsky et al., 2013). Lepidoptera seem to be especially prone to unexpected range shift responses, though it is not always clear why. For example, across studies between 22 - 33% of Lepidoptera species shifted their ranges downslope rather than upslope as expected (Chen et al., 2009; Franco et al., 2006; Konvicka et al., 2003; Lenoir et al., 2010; McCain & Garfinkel, 2021; Wilson et al., 2005). One possibility is that Lepidoptera are tracking non-temperature related variables in response to climate change such as precipitation. Precipitation is likely to be particularly important for Lepidoptera owing to their reliance on host plants (which require certain amounts of water for their own growth) for development through their juvenile stages and adult diet (Lancaster, 2020). In addition, Lepidoptera are ectothermic insects which breathe through spiracles and simultaneously lose water (respiratory water loss), meaning that preserving water balance is particularly difficult in dry environments and is especially important for maintaining homeostasis (Chown, 2002; Jogar et al., 2004).

While the magnitude and direction of local climate velocity might shape range shift responses and provide a potential explanation for longitudinal range shifts, species also differ in their intrinsic capabilities to be pushed, pulled, or resist shifting in response to climate change. In particular, the climates in which species evolved over long timescales have the potential to shape contemporary range shift responses under recent climate change. For example, species traits, such as their environmental niche breadths (e.g. range of climatic temperatures they inhabit) and dispersal capability, might also influence their range shift responses (Hällfors et al., 2023; Herrera et al., 2018; Pöyry et al., 2009). Species with broad environmental thermal niche breadths might not shift their ranges as quickly as those with a narrow thermal niche breadth (Hällfors et al., 2023; Herrera et al., 2018). This is because species with broad environmental niche breadths are likely to have broad climatic tolerances or be highly plastic to allow survival across a broad range of climatic conditions (da Silva et al., 2020; da Silva et al., 2019). Likewise, dispersal capability might govern the ability of different species to shift in response to changing climatic change. For example, wingspan, an approximation for dispersal capability in Lepidoptera, is positively associated with the magnitude of range shift response (Pöyry et al., 2009).

Lepidopteran systems are well-poised to address questions regarding the prevalence and potential mechanisms of longitudinal range shift responses and the role of local climate velocity. This taxonomic group has a proclivity towards non-typical range shifts (Lenoir et al., 2010), and is highly sensitive to changes in temperature and moisture (Cayton et al., 2015; Diamond et al., 2014; Hällfors et al., 2023; Hordley et al., 2023). Here, we take advantage of fine scale spatio-temporal monitoring of butterflies in the Midwestern United States representing 18 years of data across a spatial extent spanning over 116,000 km^2^ to explore these questions. We compute vectors of centroid shift responses (latitudinal and longitudinal components) for 88 species and link these with local climate change velocities based on temperature and precipitation. We examine whether local climate velocities can aid in explaining non-poleward shifts among species in the butterfly assemblage, and whether these local climate velocities are better predictors of the shift response than mean climate change velocity species experience throughout their entire range. We also explored how dispersal capability (wingspan) and thermal niche breadth influences range shifts, while examining the role of shared evolutionary history in mediating the shift response. In general, comparative studies of contemporary range shift responses show overall low, but quite variable, roles for evolutionary history in mediating the shift response (Diamond 2018). Previously unaccounted variation in the magnitude and direction of local climate velocities along with species niche and dispersal traits could provide an explanation for this outcome, and might therefore increase the degree to which range shift responses are explained by evolutionary history.

## Methods

### Centroid shifts

We examined range shifts in 88 species of North American butterfly species by examining changes in abundances across 146 fixed transects between the years 2000 - 2017. Transect monitoring was conducted by citizen scientists that took part in the Ohio Lepidopterists Monitoring Program within a strict range of environmental conditions on a weekly basis (https://www.ohiolepidopterists.org/). We only examined range shifts in species geographic ranges that had over 80 observations across the 18 year test period, and species that had range shift data for at least a 10 year period.

We used abundance-weighted methods to assess each species’ range shifts (Ash et al., 2017, Huang et al., 2017; 2023). We calculated the abundance-weighted geographic centroid of each species each year between 2000 and 2017 within the Ohio Lepidoptera monitoring scheme to create centroid shift vectors for each species. To account for potential sampling error we bootstrapped species occurrence records by resampling 80% of each species annual records with replacement 10,000 times to estimate each species’ mean geographic centroid each year of the survey. Centroid shift vectors have been shown to be an efficient and accurate way to assess fine scale changes in species distribution, especially over relatively short time scales (Ash et al., 2017; Huang et al., 2017, 2023).

We used linear models to calculate the velocity at which each species shifted their latitudinal and longitudinal centroids across space. The range shift velocities have components of both rate and direction. The slope of each model was extracted to estimate range shift velocity. Degrees latitude were converted to kilometres by multiplying latitude by 84 (the mean degrees to latitude conversion across Ohio’s latitudinal span), and degrees longitude were converted to kilometres by multiplying degrees longitude by 111 (same longitudinal conversion across the globe). We calculated velocity of centroid shift as per Huang et al. 2017 equation 2,

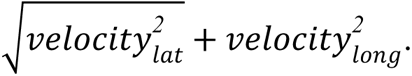

Using the slopes of the latitudinal and longitudinal centroid shifts we estimated the start and end points for each species centroid shift within Ohio to visualise range shift vectors. We also estimated the standard error associated with each species geographic range shift and accounted for this error within our models when examining the drivers of species range shifts (see details below). The R package geosphere (Hijmans et al., 2017) was used to calculate the distance (km) each species centroid shifted and the direction they shifted (bearing degrees). We plotted the rate and direction of species range shifts using a rose wind diagram from the package clifro (Seers & Shears, 2015).

### Climate change velocity

We calculated the climate change velocities that each butterfly species experiences at their modelled starting (2000) and ending (2017) centroid within Ohio as well as the mean climate change velocity species experience throughout their entire geographic distribution using the vocc (Brito-Morales et al., 2018) package. Climate change velocities are calculated as the ratio of temperature change over time (temporal trend), to temperature change over space (spatial trend) based on a 3 x 3 cell neighbourhood (Loarie et al., 2009), where climate change velocity = spatial trend/temporal trend. We estimated the velocity at which the warmest temperatures in July (hottest month of the year), coldest temperatures in January (coldest month of the year), and mean precipitation received in April (important month for plant growth and butterfly development), changed from 2000 - 2017, matching the period of time that we analyse butterfly range shifts. We also divided Ohio into quadrants and estimated the climate change velocity within each quadrant.

### Niche and dispersal traits

To assess how niche traits might be influencing species range shifts, we extracted occurrence data from the Global Biodiversity Information Facility (GBIF) for each species (list of doi’s for each species range data listed in Supplementary Table 1). We computed species distribution polygons for each species using the rangeBuilder (Davis Rabosky et al., 2016) and geosphere (Hijmans et al., 2017) packages. Using the raster (Hijmans et al., 2015) package, we estimated the maximum and minimum environmental temperatures species experience within their range polygons at a 30-arc-second resolution to calculate the environmental thermal niche breadth of each species.

As different parts of species ranges have different range shift responses to climate change (Comte et al., 2014), we calculated an index of each species geographic range position relative to Ohio. We did this using the minimum and maximum latitudes of each species range polygon to calculate a latitudinal range for each species. We then estimated the quantile of this range that overlapped the latitudinal midpoint of the monitoring scheme. Values range from 0 to 1, with values closer to 1 indicating the monitoring scheme overlaps the species’ range towards its poleward edge and values closer to 0 overlapping toward its equatorward edge.

We estimated dispersal capability based on wingspan (in cm) (Daniels, 2004). Minimal and maximal wingspan values from one forewing tip to another were available for the majority of butterfly species, so we tested the predictive ability of both sets of wingspan estimates.

### Statistical Analysis

Statistical analyses were performed in the R version 4.2.1 (R Core Team, 2023). To test the drivers of butterfly range shift responses, we used the ‘rma.mv’ function in the metafor package (Viechtbauer, 2010) so that we could account for the error in the rate of centroid shift we estimated when calculating species shift vectors, account for phylogenetic effects, and conduct hypothesis testing.

We used a recently published phylogeny of the butterflies of North America which was constructed using 13 common markers (1 mitochondrial and 12 nuclear) (Earl et al., 2021) to account for evolutionary history in our models. The phylogeny included 70/88 species that we had centroid shift data for. We examined if there was phylogenetic signal in rate of range shift, wingspan, and thermal niche breadth by calculating Pagel’s 𝜆 using the phytools package (Revell, 2012). We also examined if there was phylogenetic signal in the direction species shifted their ranges by computing discrete character evolution models using the fitDiscrete function to estimate Pagel’s 𝜆 with the package geiger (Harmon et al., 2015). We computed equal-rate (ER), symmetric transition (SYM) and all rate different (ARD) evolutionary models with a lambda transformation and then compared them using Akaike Information Criterion (AIC) and likelihood ratio tests.

We structured our models whereby species rate of range shift was the response variable, and the standard error associated with each species rate of range shift was accounted for. We included local climate change velocity (temperature and precipitation) and mean climate change velocity (temperature and precipitation) throughout species ranges in the same models, however, to avoid introducing multicollinearity into our models, we did not include maximum and minimum temperature velocity in the same model as those models had variance inflation factors (VIFs) over 2.5 (Johnston et al., 2018). Similarly, thermal niche breadth and latitudinal position relative to Ohio were correlated (r^2^ = 0.64), and models that included both variables had VIFs over 2.5. We elected to include thermal niche breadth in our models as our main hypotheses were focused on this trait; we excluded latitudinal position relative to Ohio to avoid collinearity issues. All models included species and a phylogenetic correlation matrix as random factors to account for shared evolutionary history. We used a maximum likelihood modelling approach in all models. Using model reduction (only variables linked to key hypotheses were included in the full models) and AIC comparison, we examined which models best explained variation in species range shifts. We used likelihood ratio tests for significance testing of the variables in the best-fitting models.

In a separate model, which included the predictor variables from the model with the lowest AIC value, we examined the relationship between rate of range shift and wingspan as a proxy for dispersal capability. Wingspan was analysed separately because wingspan data was only available for a subset of the species in the dataset (n = 66).

We also examined if the rate that species shifted their centroids depended on the direction that butterflies were shifting their centroids, in a model that included rate of centroid shift (and its associated standard error) as a response variable, and direction as a categorical variable (N, S, E, W, NE, NW, SE, SW). Furthermore, to understand if there is a relationship between climate change velocity (local and whole range temperature and precipitation) or wingspan on the direction species shifted their centroids, we conducted separate linear models, where climate change velocity or wingspan were included as continuous response variables and direction was included as a categorical predictor variable (as a categorical variable cannot be used as a response variable).

To test if butterflies were consistently moving either towards or away from regions of rapid climate change, we used separate linear models to examine if climate change velocity (local maximum environmental temperature and precipitation velocity) at the start of a species range shift vectors predicted climate change velocity at the end of species range shift vectors. We tested the significance of the relationship between climate change velocities at the start and end of species centroid shift vectors by using likelihood test ratios.

## Results

Butterflies shifted their centroids at an average of 4.87 km per year in all directions (Figure 1 & Supplementary Table 1 & 2), but, the vast majority of species moved in a westerly (including north-west and south-west) direction (Figure 1 & 2; Supplementary Table 2)). Centroid shift velocities ranged from 0.2 to 25.7 km per year. While 40 butterfly species significantly shifted their centroids, the remaining 48 species did not significantly shift their centroids (Supplementary Table 1).

**Figure 1.**
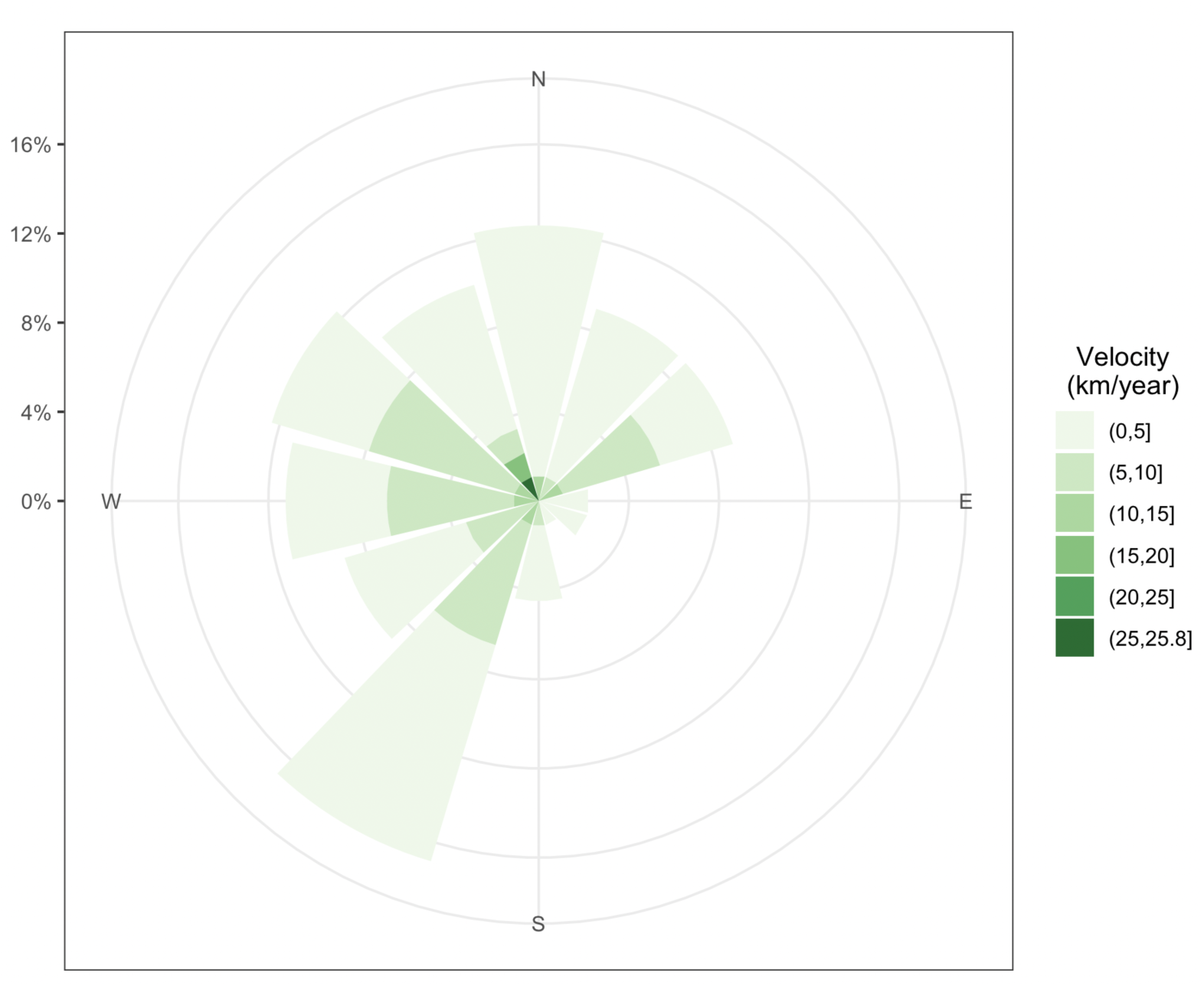
Rose wind diagram showing the velocity of butterfly centroid shifts between 2000 - 2017 in Ohio USA. Colour indicates centroid shift velocity and size indicates the proportion of the sample that shifted at a particular rate and direction.

### Phylogenetic signal

There was high phylogenetic signal in species rate of centroid shift (𝝀 = 0.96, P = 0.013), meaning that species that are more closely related have more similar rates of range shift than distantly related species (Figure 2). There was also high phylogenetic signal in lower (𝝀 = 0.98, P < 0.05) and upper wingspan length (𝝀 = 1.0, P < 0.05). However, we found low phylogenetic signal in the environmental thermal niches species inhabit (𝝀 = < 0.01, P = 1).

**Figure 2.**
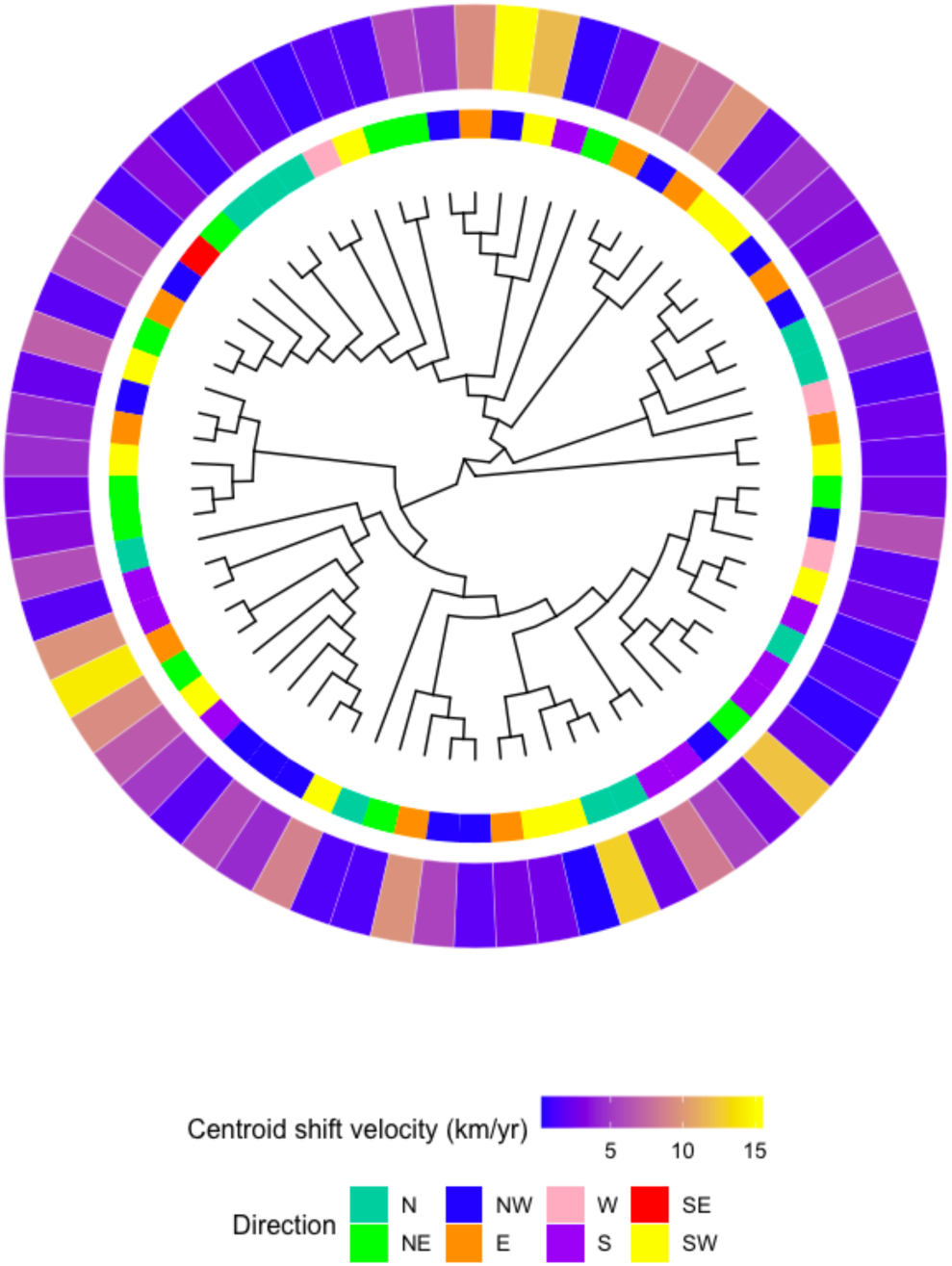
Pruned ultraconserved element North American butterfly phylogeny (Earl et al. 2021). Tip colours correspond to the direction that butterflies shifted their centroid within Ohio and the centroid shift velocity (km yr^-1^) of butterflies between 2000 - 2017 (n = 70).

To determine if closely related species were more or less likely to shift in similar directions, we computed and compared discrete character evolution models with different rates of state change. The equal rates model had the lowest AIC value (ER = 296.72, SYM = 327.92, ARD = 375.78); however, likelihood ratio tests indicated that no models were significantly different from each other. The equal rate discrete character evolution model had a Pagel’s 𝜆 value of 0.79 (SYM = 0.89 , ARD = 0.71), suggesting that closely related species are more likely to shift their geographic ranges in similar directions (Figure 2).

### What are the drivers of butterfly range shift responses?

We found that a model that included local maximum temperature velocity and local precipitation velocity best explained variation in butterfly centroid shifts (Figures 3A & B; see model AIC comparisons in Supplementary Table 4). Rate of centroid shift was positively correlated with both maximum temperature (estimate = 29.34 ± 3.94, Z = 7.45, 𝝌^2^ = 41.04, P < 0.001) and precipitation velocity (estimate = 0.39 ± 0.06, Z = 6.69, 𝝌^2^ = 36.75, P < 0.001) (Figure 3A & B). A model that included thermal niche breadth as well as local maximum temperature and precipitation velocity had the second lowest AIC values which had a delta AIC of less than 2 (indicating the models explain a similar amount of variation in rate of centroid shift). However, when we conducted a likelihood ratio test, thermal niche breadth was not significantly associated with rate of centroid shift (𝝌^2^ = 1.14, df = 1, P = 0.286). In a separate model, we included wingspan data from a reduced sample where wingspan data was available. Wingspan was not associated with rate of centroid shift (𝝌^2^ = 0.86, df = 1, P = 0.86).

**Figure 3.**
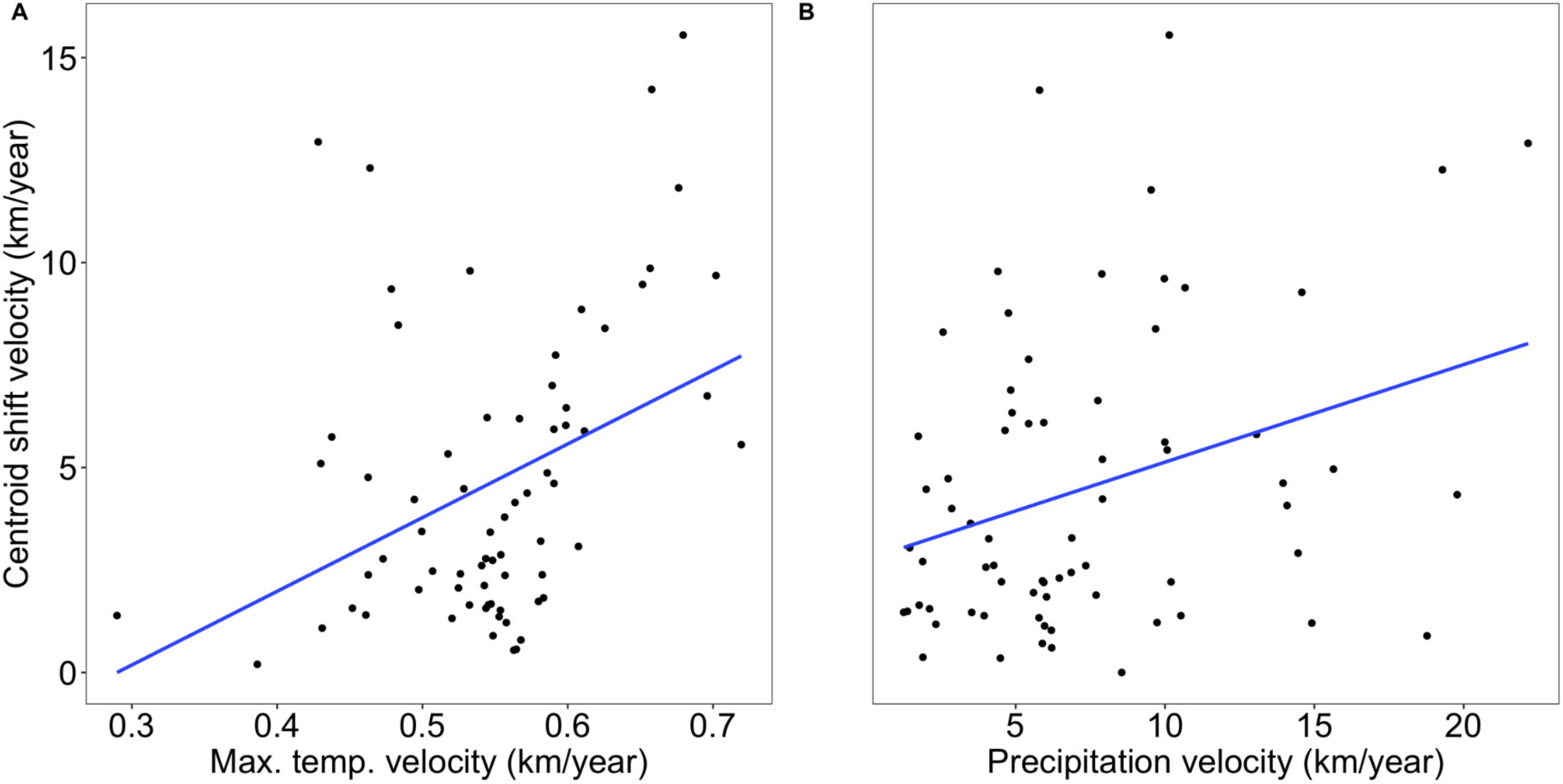
Relationship between rate of centroid shift and A) local maximum temperature velocity, and B) local precipitation velocity.

### Are butterflies shifting their ranges in predictable ways?

The speed at which species shifted their centroid also depended on the direction that species shifted their ranges (𝝌^2^ = 20.44, df = 1, P = 0.005). Butterflies shifted their centroids fastest in westerly directions (northwest, west and southwest) (Figure 1 & 2). Only one species shifted their centroid in a southeasterly direction, which is also the quadrant of Ohio that was warming at the fastest velocity (Figures 1 & 4; Supplementary Table 3). Many species shifted their centroids in a northwesterly or southwesterly direction towards regions that were increasing in precipitation velocity (getting wetter).

**Figure 4.**
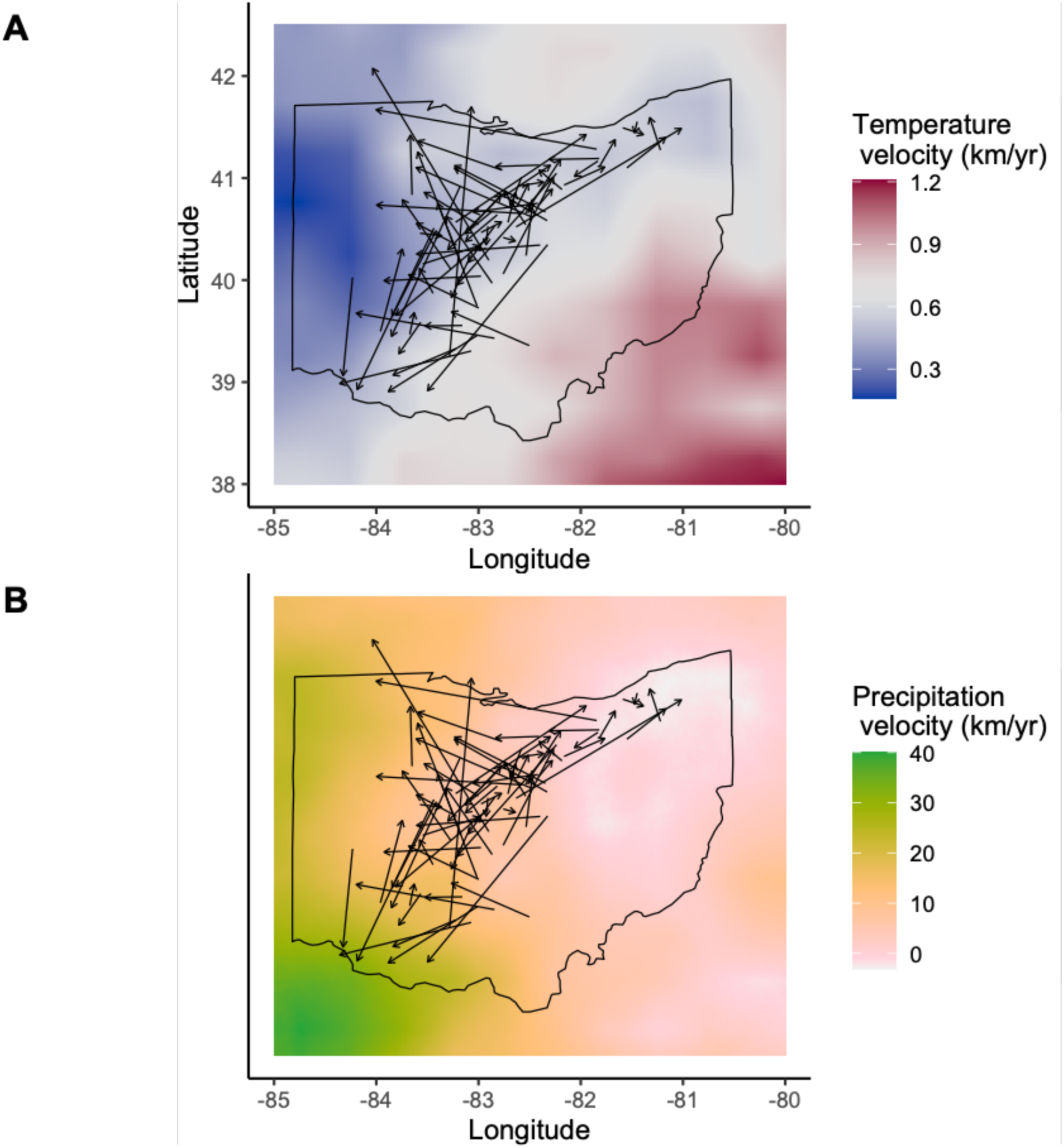
Estimated butterfly species range shift vectors from the estimated start and end centroids from each species linear mixed effect models between 2000 and 2017.

We also examined if species shifted their centroids into regions that are changing more slowly (i.e. lower climate change velocities) by examining the relationship between temperature and precipitation velocity at the start and end of species range shift vectors. There was no significant relationship between the local temperature velocity that species experienced at their starting and ending centroid (𝝌^2^ = 0.039, df = 1, P = 0.843). However, there was a positive relationship between local starting precipitation velocity and end of range shift precipitation velocity (𝝌^2^ = 14.97, df = 1, P < 0.001; R^2^ = 0.19, Figure 5). This suggests that many species appear to be tracking precipitation velocities. Climate change velocity (local and whole range temperature and precipitation) and species wingspan did not influence the direction butterflies shifted their centroid (P > 0.005).

**Figure 5.**
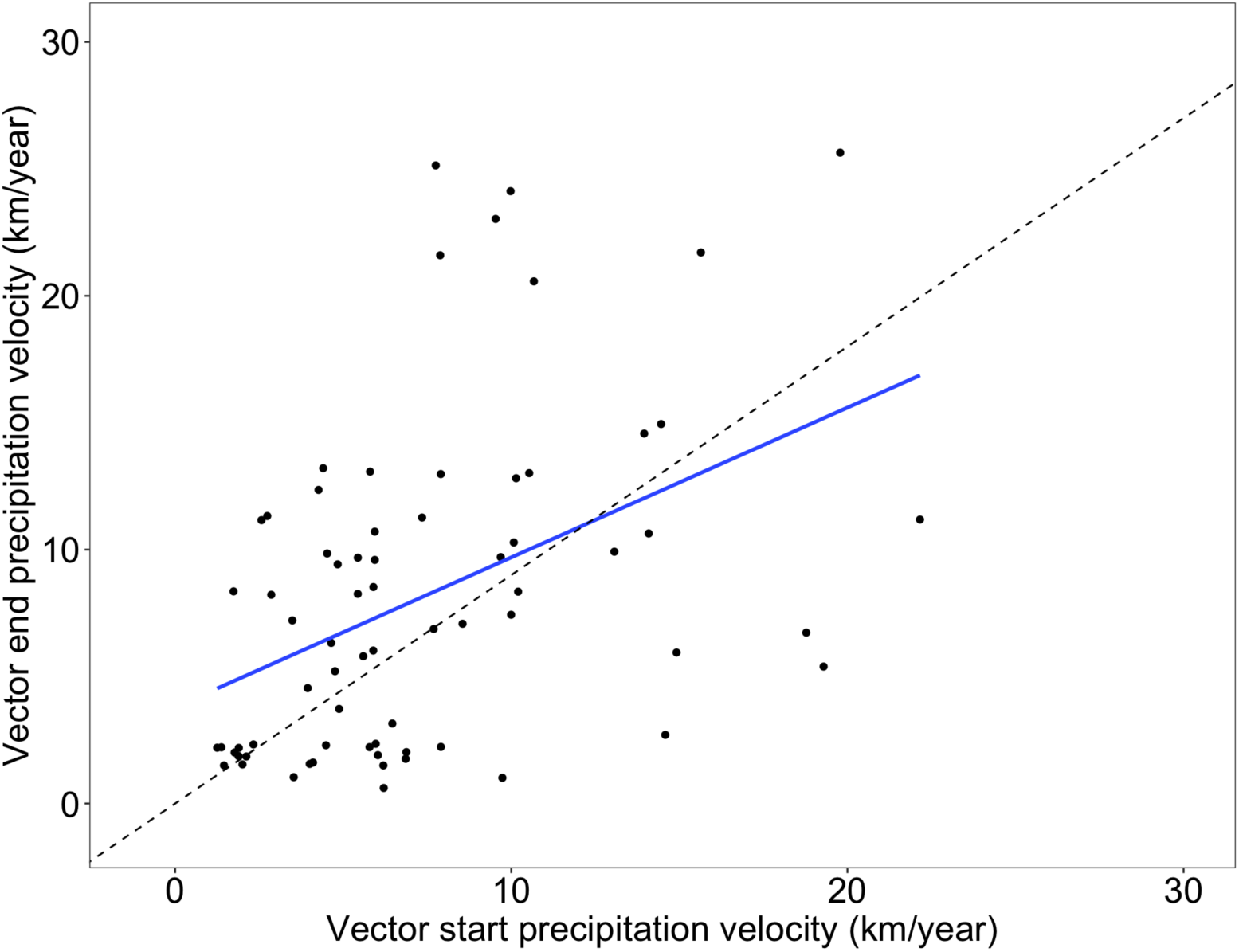
Relationship between the precipitation velocity species experienced at the start of the centroid shift vector and the end of their centroid shift vector. Points above the dotted line indicate butterflies that shifted their ranges into environments that are becoming wetter at a faster rate than the precipitation velocity at their starting centroid. Points below the dotted line represent species that shifted their centroids into areas that are increasing precipitation at a slower rate than their starting position. While some regions in Ohio are getting more arid (precipitation velocities below 0), the start and end of centroid shift points did not overlap with these areas.

## Discussion

Until recently, most studies have examined poleward and upslope shifts in species ranges to avoid warming climates (Chen et al., 2009, 2011; Lenoir & Svenning, 2015; Parmesan & Yohe, 2003; VanDerWal et al., 2013). By ignoring longitudinal range shift components, it is possible that we are underestimating species range shift responses and are missing important directional information. By conducting multidirectional centroid shift analyses (considering latitudinal and longitudinal shifts over an 18 year period), we found that butterflies in the Midwestern United States are shifting their centroids at different rates and in all directions, except towards regions that are warming the fastest. This study, along with a few other recent publications (Hordley et al., 2023; Huang et al., 2017; Lenoir et al., 2010, 2020; Pinsky et al., 2013; VanDerWal et al., 2013), challenges the notion that species are solely shifting their ranges poleward and upslope to avoid warming habitats and highlights that changes in precipitation regimes also play a major role in shaping species range shift responses.

We found strong longitudinal components to butterfly species centroid shift responses. A majority of butterflies shifted their centroids in a westerly (west, northwest, southwest) direction (Figure 1, Supplementary Table 2). The western quadrants of Ohio are warming at slower rates than the eastern quadrants, and are also becoming wetter at faster rates than the eastern quadrants (Figure 3; Supplementary Table 3). This suggests that butterflies are moving away from environments that are heating up quickly and that some species are moving into regions that are getting wetter. Only one species of butterfly (*Polites mystic*) shifted their centroid in a southeasterly direction (towards the quadrant that is warming the fastest), and the rate that they shifted their centroid was slow (southeast shift velocity = 1.4 km yr^-1^) compared to butterflies that shifted in a westerly direction (towards regions that are warming at a slower rate and becoming wetter at a faster rate) (west shift mean velocity = 6.9 km yr^-1^) (Figure 1; Supplementary Table 2). Overall, the rate that butterflies shifted their centroids was strongly influenced by the velocity of local climate change (maximum temperature and precipitation) (Figure 2, 3 & 4).

Our finding that centroid shifts (direction and rate) depend on both temperature and precipitation velocity aligns well with other studies. For example, changes in moth distributions are associated with temperature and precipitation, whereby leading edge populations are shifting north with changes in temperature, but centroids are shifting northwest with changes in precipitation (Hordley et al., 2023). While the local temperature velocity at the start of species shift vectors did not predict the thermal velocity environment they moved into, butterflies tended to track precipitation velocities over the 18 year period (Figure 5). Interestingly, however, some species that started in locations with high precipitation velocities (likely species that need wet conditions) moved into even high precipitation velocity locations, where species that started in low precipitation velocity areas don’t seem to track precipitation velocities as closely (Figure 5). Changing precipitation regimes are likely to play a major role in reshaping Lepidoptera distributions as they rely on plants for development and diet, and plant richness and abundance is correlated with precipitation, especially in dry habitats (Korell et al., 2021). Thus, butterflies might not just be following changes in precipitation, but could be following host plant distributions and abundance, potentially also explaining longitudinal components of species range shifts. Or butterflies could be moving to environments that are increasing in precipitation to avoid desiccating environments which are associated with high temperatures and low precipitation.

Combined changes in temperature and precipitation influence desiccation stress or drying power of the air (vapour pressure deficit), where desiccation stress is highest in warm and dry environments (White et al., 2007). Our finding that butterflies are moving away from regions that are becoming warmer at a rapid rate, and towards regions that are becoming wetter at a faster rate suggest that butterflies could be escaping regions of higher desiccation stress. Insects are particularly prone to respiratory water loss as they breathe through spiracles which simultaneously allow oxygen into the tracheal system, and release carbon dioxide and water (Chown, 2002). Desiccation stress in the environment is known to shape the geographic distributions and desiccation resistance of insects such as *Drosophila* (Kellermann et al., 2012). Thus, changes in precipitation regimes are likely to induce range shift responses in species that are not adapted to desiccating environments, and likely contribute towards longitudinal components of species range shifts.

Longitudinal components of species range shifts may also be explained by the finding that local climate change velocity has a greater influence on species range shifts than the mean climate change velocity throughout a species range. Climate change velocity is highly variable across space, and it’s likely that butterflies within local environments are responding to direct changes in local climate, rather than responding to change in climate as a species with a unified response. This is important because it implies that populations within species might be shifting their geographic ranges at multiple rates and directions depending on the changes in local climates they experience (Lenoir et al., 2020; Lenoir & Svenning, 2015; Pinsky et al., 2013). Previous studies have found that range shift dynamics differ in centroid, leading and trailing edge populations (Comte et al., 2014; Hordley et al., 2023; Lenoir et al., 2020), supporting this idea. In addition, it is acknowledged that species range shifts are sensitive to variation in their initial/historic geographic distributions, and thus, these initial distributions will influence the local climate change velocities that species are exposed to, and therefore must respond to (Trisos et al., 2020; Pigot et al., 2023). Indeed, impacts of species initial geographic distributions, will also impact the degree of climatic stress and timing of climate change events that stimulate different species (and populations within species initial ranges) to shift their ranges. Future research should seek to examine how climate change velocity is correlated with multiple populations within species to examine how population level range shifts interact to influence whole species range shifts.

Thermal niche breadth did not explain significant variation in species range shift responses. If thermal niche breadths are representative of species thermal tolerance breadths, we would have expected that species with narrow thermal niches would shift their ranges more quickly to track climates they can survive in. However, adaptation to local environments might mean that thermal niche breadths of an entire species do not match the thermal tolerance breadths of individuals at particular locations (Sunday et al., 2015). We also found no evidence to support that wingspan, which is often used as a proxy for dispersal capability, influences rate of range shift. Previous studies provide mixed support for whether wingspan is correlated with range shift responses (Kharouba et al., 2009; Pöyry et al., 2009). However, in the current study, when local climate change velocity and evolutionary history are accounted for, wingspan and thermal niche breadth did not play an important role in predicting range shift responses in butterflies in the Midwestern United States.

While we found that local climate change velocity plays an important role in determining species range shift responses, we also found strong phylogenetic signal in species range shift responses (rate and direction). Across taxonomic groups, there is a great deal of variation in phylogenetic signal in range shift responses, however, overall, phylogenetic signal tends to be low (Diamond, 2018). This could mean that range shift responses might be a relatively labile trait, or it could indicate that phylogenetic signal is being underestimated due to missing information on the longitudinal components of species range shifts. Similarly, very few studies have examined phylogenetic signal in range shift direction, especially while considering longitudinal components of species range shifts.

## Conclusions

Butterflies in the Midwestern United States are shifting their centroids in all directions. They are moving away from areas that are warming at the most rapid rates and many species are tracking precipitation velocity. Local climate change velocity better predicted rates of centroid shift than the climate change velocity species experience throughout their entire ranges, suggesting that populations within species might be shifting their ranges in multiple ways. These findings highlight that range shift responses to climate change are complex and vary depending on the climate change velocity in local regions and also on species evolutionary histories.

## Acknowledgements

We would like to thank Jerry Wiedmann and the many volunteer citizen scientists of the Ohio Lepidopterists who dedicated their time to monitor butterflies.

## Funding

This work was supported by the National Science Foundation (DEB-1845126 to Sarah E. Diamond).

## Data availability

Data will be archived on dryad upon manuscript acceptance.

## Notes

### Competing Interest Statement

The authors have declared no competing interest.

### Summary of Updates

We have amended the manuscript in response to reviewer suggestions. We have also added the GBIF download dois that we used to make each species range polygon.

