## Supplementary tables for "Local climate change velocities explain multidirectional range shifts in a North American butterfly assemblage"

**Supplementary Material**

**Supplementary Table 1.** Summary of centroid shift responses for butterflies in Ohio USA.

| Species | Year span | Centroid shift velocity (km/yr) | Significant centroid shift | Bearing (clockwise) | Direction | Family | GBIF doi for range trait calculations |
|---|---|---|---|---|---|---|---|
| *Achalarus lyciades* | 17 | 9.86 | yes | 273.88 | W | Hesperiidae | 10.15468/dl.4e0pps |
| *Amblyscirtes vialis* | 16 | 11.82 | yes | 212.02 | SW | Hesperiidae | 10.15468/dl.h2vnbo |
| *Ancyloxypha numitor* | 18 | 2.74 | yes | 59.73 | NE | Hesperiidae | 10.15468/dl.zrd4o8 |
| *Anthocharis midea* | 17 | 4.48 | yes | 241.25 | SW | Pieridae | 10.15468/dl.sqjboy |
| *Asterocampa celtis* | 18 | 8.48 | yes | 199.08 | S | Nymphalidae | 10.15468/dl.ui902a |
| *Asterocampa clyton* | 17 | 5.33 | no | 201.47 | S | Nymphalidae | 10.15468/dl.2rncnn |
| *Atalopedes campestris* | 18 | 1.08 | no | 8.98 | N | Hesperiidae | 10.15468/dl.d78gjb |
| *Atrytone logan* | 18 | 9.67 | yes | 284.55 | W | Hesperiidae | 10.15468/dl.nnrbz1 |
| *Atrytonopsis hianna* | 14 | 15.55 | yes | 335.09 | NW | Hesperiidae | 10.15468/dl.tiz9v0 |
| *Battus philenor* | 18 | 2.38 | no | 268.81 | W | Papilionidae | 10.15468/dl.iotqlo |
| *Boloria bellona* | 18 | 2.37 | no | 5.38 | N | Nymphalidae | 10.15468/dl.twqwbp |
| *Boloria selene* | 17 | 12.94 | yes | 3.9 | N | Nymphalidae | 10.15468/dl.a0bqk9 |
| *Callophrys gryneus* | 17 | 5.1 | yes | 184.27 | S | Lycaenidae | 10.15468/dl.atqkfr |
| *Callophrys henrici* | 17 | 9.8 | yes | 199.88 | S | Lycaenidae | 10.15468/dl.u8hazr |
| *Calycopis cecrops* | 17 | 6.75 | yes | 230.56 | SW | Lycaenidae | 10.15468/dl.iinubp |
| *Celastrina ladon* | 18 | 1.57 | yes | 196.33 | S | Lycaenidae | 10.15468/dl.7545oq |
| *Cercyonis pegala* | 18 | 1.22 | no | 32.78 | NE | Nymphalidae | 10.15468/dl.p4gzqb |
| *Chlosyne nycteis* | 18 | 2.38 | no | 199.3 | S | Nymphalidae | 10.15468/dl.hzucag |
| *Colias cesonia* | 15 | 10.32 | no | 311.32 | NW | Pieridae | 10.15468/dl.evqjgr |
| *Colias eurytheme* | 18 | 4.15 | yes | 259.5 | W | Pieridae | 10.15468/dl.niy7ez |
| *Colias philodice* | 18 | 2.12 | yes | 334.71 | NW | Pieridae | 10.15468/dl.vxj3s9 |
| *Cyllopsis gemma* | 16 | 9.68 | yes | 251.9 | W | Nymphalidae | 10.15468/dl.fku4tu |
| *Danaus plexippus* | 18 | 1.32 | no | 20.23 | N | Nymphalidae | 10.15468/dl.1hxwg4 |
| *Enodia anthedon* | 18 | 2.95 | yes | 36.72 | NE | Nymphalidae | 10.15468/dl.0i1vei |
| *Epargyreus clarus* | 18 | 2.02 | no | 222.42 | SW | Hesperiidae | 10.15468/dl.epaybx |
| *Erynnis baptisiae* | 18 | 3.2 | no | 268.42 | W | Hesperiidae | 10.15468/dl.xtnmdu |
| *Erynnis brizo* | 17 | 5.93 | no | 343.55 | N | Hesperiidae | 10.15468/dl.gjpzit |
| *Erynnis horatius* | 18 | 3.79 | yes | 307.98 | NW | Hesperiidae | 10.15468/dl.tvp2h5 |
| *Erynnis icelus* | 17 | 4.87 | no | 331.85 | NW | Hesperiidae | 10.15468/dl.ewdyaw |
| *Erynnis juvenalis* | 18 | 4.61 | no | 216.89 | SW | Hesperiidae | 10.15468/dl.2clqkh |
| *Euphydryas phaeton* | 17 | 12.31 | no | 52.31 | NE | Nymphalidae | 10.15468/dl.blxziv |
| *Euphyes conspicuus* | 15 | 4.51 | no | 241.86 | SW | Hesperiidae | 10.15468/dl.dbrvrj |
| *Euphyes dukesi* | 12 | 5.75 | no | 67.4 | NE | Hesperiidae | 10.15468/dl.j7kqs3 |
| *Euphyes vestris* | 18 | 1.4 | no | 56.16 | NE | Hesperiidae | 10.15468/dl.sapng5 |
| *Euptoieta claudia* | 17 | 0.2 | no | 211.59 | SW | Nymphalidae | 10.15468/dl.znhr7q |
| *Eurema lisa* | 18 | 4.6 | yes | 12.25 | N | Pieridae | 10.15468/dl.epw7ia |
| *Eurema nicippe* | 18 | 9.74 | yes | 55.83 | NE | Pieridae | 10.15468/dl.yxcv4c |
| *Eurytides marcellus* | 18 | 2.06 | no | 208.32 | SW | Papilionidae | 10.15468/dl.e3jomb |
| *Everes comyntas* | 18 | 2.59 | yes | 253.05 | W | Lycaenidae | 10.15468/dl.b5y1ym |
| *Feniseca tarquinius* | 17 | 6.03 | yes | 21.75 | N | Lycaenidae | 10.15468/dl.vseywl |
| *Hermeuptychia sosybius* | 17 | 5.56 | yes | 299.11 | NW | Nymphalidae | 10.15468/dl.yjzbeh |
| *Hesperia leonardus* | 17 | 3.08 | no | 358.81 | N | Hesperiidae | 10.15468/dl.3w6ikj |
| *Hylephila phyleus* | 18 | 3.42 | no | 26.34 | NE | Hesperiidae | 10.15468/dl.ulcaf7 |

| | | | | | | | |
|---|---|---|---|---|---|---|---|
| *Junonia coenia* | 18 | 2.77 | no | 304.9 | NW | Nymphalidae | 10.15468/dl.fz2dcq |
| *Libytheana carinenta* | 18 | 4.59 | yes | 204.07 | SW | Nymphalidae | 10.15468/dl.u1cx61 |
| *Limenitis archippus* | 18 | 2.78 | no | 267.26 | W | Nymphalidae | 10.15468/dl.olfgel |
| *Limenitis arthemis* | 18 | 2.47 | yes | 208.96 | SW | Nymphalidae | 10.15468/dl.bomx99 |
| *Lycaena hyllus* | 18 | 9.35 | yes | 48.78 | NE | Lycaenidae | 10.15468/dl.nc7pdj |
| *Lycaena phlaeas* | 18 | 14.22 | yes | 282.66 | W | Lycaenidae | 10.15468/dl.xejfik |
| *Megisto cymela* | 18 | 1.73 | no | 318.13 | NW | Nymphalidae | 10.15468/dl.pdfodo |
| *Nastra lherminier* | 17 | 9.47 | yes | 282.8 | W | Hesperiidae | 10.15468/dl.rs312t |
| *Nymphalis antiopa* | 18 | 1.52 | no | 10.73 | N | Nymphalidae | 10.15468/dl.vurikp |
| *Nymphalis milberti* | 17 | 6.56 | no | 61.69 | NE | Nymphalidae | 10.15468/dl.tretdu |
| *Papilio cresphontes* | 18 | 1.7 | yes | 164.7 | S | Papilionidae | 10.15468/dl.jxa7go |
| *Papilio glaucus* | 18 | 0.99 | no | 348.18 | N | Papilionidae | 10.15468/dl.benwrj |
| *Papilio troilus* | 18 | 3.05 | no | 309.86 | NW | Papilionidae | 10.15468/dl.vrfxv4 |
| *Phoebis sennae* | 18 | 7 | yes | 222.03 | SW | Pieridae | 10.15468/dl.u4fwa0 |
| *Pholisora catullus* | 18 | 1.39 | no | 98.47 | E | Hesperiidae | 10.15468/dl.zgdwyb |
| *Phyciodes tharos* | 18 | 0.55 | no | 190.39 | S | Nymphalidae | 10.15468/dl.zahj7i |
| *Pieris rapae* | 18 | 3.44 | yes | 48.86 | NE | Pieridae | 10.15468/dl.zhpost |
| *Pieris virginiensis* | 18 | 2.87 | no | 43.92 | NE | Pieridae | 10.15468/dl.dp5w9g |
| *Poanes hobomok* | 18 | 1.65 | no | 23.74 | NE | Hesperiidae | 10.15468/dl.i2wzlo |
| *Poanes viator* | 17 | 1.82 | yes | 346.13 | N | Hesperiidae | 10.15468/dl.xrubk8 |
| *Poanes zabulon* | 18 | 0.79 | no | 110.2 | E | Hesperiidae | 10.15468/dl.511ani |
| *Polites mystic* | 18 | 1.36 | no | 113.88 | SE | Hesperiidae | 10.15468/dl.naxqyd |
| *Polites origenes* | 18 | 4.76 | no | 301.49 | NW | Hesperiidae | 10.15468/dl.ri03f3 |
| *Polites peckius* | 18 | 0.87 | no | 354.53 | N | Hesperiidae | 10.15468/dl.mksmdk |
| *Polites themistocles* | 18 | 6.19 | yes | 266.2 | W | Hesperiidae | 10.15468/dl.goiprp |
| *Polygonia comma* | 18 | 0.9 | no | 184.35 | S | Nymphalidae | 10.15468/dl.zo2rud |
| *Polygonia interrogationis* | 18 | 2.41 | no | 222.17 | SW | Nymphalidae | 10.15468/dl.nsb9po |
| *Pompeius verna* | 18 | 1.67 | no | 229.16 | SW | Hesperiidae | 10.15468/dl.59g8tc |
| *Pyrgus communis* | 18 | 4.22 | no | 11.62 | N | Hesperiidae | 10.15468/dl.lwdgi1 |
| *Satyrium calanus* | 18 | 4.38 | yes | 332.04 | NW | Lycaenidae | 10.15468/dl.nzxwnr |
| *Satyrium edwardsii* | 15 | 7.59 | yes | 288.87 | W | Lycaenidae | 10.15468/dl.1ezh9v |
| *Satyrium liparops* | 17 | 8.86 | no | 213.89 | SW | Lycaenidae | 10.15468/dl.ma7siy |
| *Satyrium titus* | 17 | 5.89 | yes | 294.59 | NW | Lycaenidae | 10.15468/dl.uyjcjb |
| *Satyrodes appalachia* | 17 | 2.27 | no | 228.03 | SW | Nymphalidae | 10.15468/dl.hhegr2 |
| *Satyrodes eurydice* | 17 | 7.44 | yes | 230.61 | SW | Nymphalidae | 10.15468/dl.cpnraj |
| *Speyeria aphrodite* | 17 | 2.13 | no | 10.86 | N | Nymphalidae | 10.15468/dl.6emqmw |
| *Speyeria cybele* | 18 | 0.75 | no | 324.85 | NW | Nymphalidae | 10.15468/dl.d0ytzn |
| *Strymon melinus* | 17 | 1.57 | no | 337.19 | NW | Lycaenidae | 10.15468/dl.wgbg0s |
| *Thorybes bathyllus* | 18 | 7.74 | yes | 299.08 | NW | Hesperiidae | 10.15468/dl.q6btaa |
| *Thorybes pylades* | 17 | 8.4 | yes | 264.18 | W | Hesperiidae | 10.15468/dl.sj3yxx |
| *Thymelicus lineola* | 18 | 0.57 | no | 190.12 | S | Hesperiidae | 10.15468/dl.efcrv1 |
| *Vanessa atalanta* | 18 | 1.65 | no | 78.84 | E | Nymphalidae | 10.15468/dl.qso4v7 |
| *Vanessa cardui* | 18 | 2.61 | no | 35.31 | NE | Nymphalidae | 10.15468/dl.idmtde |
| *Vanessa virginiensis* | 18 | 6.22 | yes | 303.66 | NW | Nymphalidae | 10.15468/dl.9eqfpy |
| *Wallengrenia egeremet* | 18 | 6.46 | yes | 306.52 | NW | Hesperiidae | 10.15468/dl.g2ic1d |

**Supplementary Table 2.** Summary of butterfly range shift responses between 2000 - 2017 in Ohio USA.

| **Direction** | **No. of species** | **Velocity km year$^{-1}$** |
|---|---|---|
| N | 14 | 3.49 |
| NE | 14 | 4.7 |
| NW | 17 | 5.1 |
| S | 10 | 3.6 |
| SE | 1 | 1.4 |
| SW | 16 | 4.6 |
| E | 3 | 1.3 |
| W | 13 | 6.9 |

**Supplementary Table 3.** Median climate change velocities (maximum environmental temperature in the hottest month (July) and precipitation in Spring (April)) within Ohio split into quadrants.

| **Quadrant** | **NW** | **NE** | **SW** | **SE** |
|---|---|---|---|---|
| Local max. temperature velocity (km/year) | 0.42 | 0.63 | 0.57 | 0.95 |
| Local precipitation velocity (km/year) | 13.11 | -0.79 | 21.15 | 3.54 |

**Supplementary Table 4.** AIC comparison of models examining which predictor variables we expect to play an important role in range shift responses explain the greatest amount of variation in species centroid shifts. Maximum and minimum temperature velocity were not included in the same models to avoid multicollinearity. Similarly, thermal niche breadth and latitudinal position relative to Ohio were correlated, and thus latitudinal position relative to Ohio (to indicate whether the group of individuals within each species were part of the central, leading or trailing edge of a species) was dropped and thermal niche breadth was kept in the models. Rate of centroid shift and its associated standard error was included as the response variable in each model and phylogeny was included as a correlation matrix. Tbreadth = thermal niche breadth; LMaxTV = local maximum temperature velocity; LMinTV = local minimum temperature velocity; LPV = local precipitation velocity; RMaxTV = species range maximum temperature velocity; RMinTV = species range minimum temperature velocity; RPV = species range precipitation velocity.

| **Predictor variables** | **df** | **AIC** | **Delta AIC** |
|---|---|---|---|
| LMaxTV + LPV | 4 | 361.813 | 0 |
| Tbreadth + LMaxTV + LPV | 5 | 362.677 | 0.864 |
| LMaxTV + LPV + RMaxTV + RPV | 6 | 364.4542 | 2.6412 |
| Tbreadth + LMaxTV + LPV + RMaxTV + RPV | 7 | 365.5717 | 3.7587 |
| Tbreadth + LMinTV + LPV + RMinTV + RPV | 7 | 389.0894 | 27.2764 |
| LMinTV + LPV + RMinTV + RPV | 6 | 390.5686 | 28.7556 |
| Tbreadth + LMaxTV + RMaxTV | 5 | 391.48 | 29.667 |
| Tbreadth + LMaxTV | 4 | 391.5941 | 29.7811 |
| LMaxTV + RMaxTV | 4 | 393.6995 | 31.8865 |
| LMaxTV | 3 | 396.559 | 34.746 |
| Tbreadth + RMinTV + RPV | 5 | 397.3147 | 35.5017 |
| LPV + RPV | 4 | 398.0187 | 36.2057 |
| Tbreadth + LPV + RPV | 5 | 398.4513 | 36.6383 |
| Tbreadth + RMinTV | 4 | 399.558 | 37.745 |
| RMinTV + RPV | 4 | 399.7929 | 37.9799 |
| Tbreadth + LMinTV + LPV | 5 | 400.1038 | 38.2908 |
| LPV | 3 | 400.8599 | 39.0469 |
| LMinTV + LPV | 4 | 402.315 | 40.502 |
| Tbreadth + RMaxTV | 4 | 403.184 | 41.371 |
| RPV | 3 | 403.5329 | 41.7199 |

| | | | |
|---|---|---|---|
| Tbreadth + RMaxTV + RPV | 5 | 404.1721 | 42.3591 |
| Tbreadth | 3 | 404.7768 | 42.9638 |
| RMaxTV + RPV | 4 | 404.7872 | 42.9742 |
| RMaxTV | 3 | 405.1536 | 43.3406 |
| Tbreadth + LMinTV | 4 | 405.3252 | 43.5122 |
| RMinTV | 3 | 407.7244 | 45.9114 |
| LMinTV | 3 | 411.0219 | 49.2089 |
